# Inferring Accumulation Times of Mitochondrial DNA Deletion Mutants from Cross-Sectional Single-Cell Data: Methodological Framework and Validation

**DOI:** 10.64898/2026.01.30.702736

**Authors:** Axel Kowald, Thomas B L Kirkwood

## Abstract

The accumulation of mitochondrial DNA (mtDNA) deletion mutants in post-mitotic cells is a hallmark of mammalian ageing and a key contributor to tissue decline in skeletal muscle and neurons. Although the occurrence of such deletions is well documented, the mechanisms by which they clonally expand to levels exceeding the threshold for respiratory chain dysfunction remain unresolved. A transcription-coupled replication model has recently been advanced, predicting that deletions affecting genes involved in a negative feedback mechanism gain a selective replication advantage. This model implies relatively short accumulation times for mutant takeover, a critical but experimentally inaccessible parameter since single-cell measurements are destructive. Here, we present a novel approach to infer such accumulation times from cross-sectional single-cell RNA sequencing (scRNAseq) data, exploiting the fact that mtDNA deletions are also reflected at the transcript level. To establish feasibility, we generated synthetic datasets using two stochastic models of the mitochondrial life cycle and used these as a gold standard. We then applied the Moran process, a classical stochastic model of birth-death dynamics, to calculate distributions of mutant accumulation times and to extract key parameters. The Moran model reproduced the distributions obtained from stochastic simulations with high fidelity, demonstrating robustness across different assumptions about mitochondrial regulation. By fitting the model to synthetic data, we successfully recovered the true values for key parameters like mutation probability, selection advantage, and the fraction of advantageous mutants. Our findings establish a methodological framework for estimating mtDNA mutant dynamics from single-cell transcriptomic data. This approach opens the way for application to large-scale datasets such as Tabula Muris Senis and Tabula Sapiens, offering new insights into the role of mtDNA deletions in ageing and age-related disease.

## Introduction

A hallmark of the ageing process is the accumulation of mitochondrial DNA (mtDNA) deletion mutants in post-mitotic cells such as neurons and skeletal muscle fibres^1-11^. Numerous single-cell and microdissection studies have shown that these deletions, though initially rare, can clonally expand within individual cells to levels well above the threshold required to impair respiratory chain function^12-15^. The resulting cytochrome c oxidase (COX) deficiency and electron transport system (ETS) abnormalities are tightly correlated with structural and functional decline of affected cells. This phenomenon has been described across species, including rodents, rhesus monkeys, and humans, establishing mitochondrial genome instability as a conserved and universal feature of mammalian ageing.

Although the occurrence of deletions is well established, the question of how they accumulate to such high intracellular levels remains unresolved. The selective forces or mechanistic processes that allow mutant genomes to outcompete wild-type mtDNA are still debated^16-21^. Early ideas emphasized the so-called “vicious cycle” hypothesis, where defective mitochondria were proposed to produce excess reactive oxygen species (ROS), thereby inducing further mutations^20,21^. However, this model fails to account for the observation that single cells typically harbour a dominant deletion rather than a mixture of multiple mutants^18^. Later explanations have included the “survival of the smallest” hypothesis^22,23^, positing that shorter genomes replicate faster, and the “survival of the slowest” model, suggesting that dysfunctional mitochondria are spared from degradation due to reduced ROS output^19^. Yet, both models face conceptual and experimental limitations. For example, genome size differences are unlikely to substantially influence replication kinetics^24^, and selective mitophagy mechanisms appear to preferentially eliminate, not protect, dysfunctional mitochondria.

A major advance came from mathematical modelling studies showing that stochastic processes alone, specifically random genetic drift under relaxed mtDNA replication, could in principle account for clonal expansion^16,17^. Neutral drift predicts that, given sufficient time and small effective mtDNA copy numbers within individual cells, some mutations arising early in life can randomly reach high frequencies. Such models reproduce the observed prevalence of COX-deficient cells in aged human muscle and brain. However, a fundamental limitation of the neutral drift hypothesis is that it only provides a good fit for long-lived species such as humans. In short-lived animals like mice or rats, the high mutation rates required to generate the experimentally observed number of COX-deficient fibres would also predict the presence of many different mutant species coexisting within the same cell, a pattern that is not seen experimentally^25^. Instead, affected cells typically harbour a single dominant deletion. This inconsistency highlights the need for additional mechanisms beyond drift to explain clonal expansion in short-lived species^26^.

To address these contradictions, new mechanistic proposals have been put forward. Kowald and Kirkwood proposed that mtDNA replication in metazoans, which is primed via transcription, creates a regulatory vulnerability^26,27^. They suggest, that under normal conditions, transcription is subject to feedback inhibition once sufficient respiratory chain subunits are produced. If a deletion removes the gene(s) involved in this feedback loop, the mutant genome escapes regulation, leading to persistently elevated transcription and replication initiation. Such mutants would thus gain a cis-acting replication advantage independent of genome size or ROS production. Consistent with this idea, surveys of deletion spectra across species have repeatedly shown that clonally expanded deletions almost always encompass the ND4 gene, and often ND5, strongly suggesting that these genes are central to the feedback control system^27^. Computational models incorporating this mechanism reproduce the low heteroplasmy observed in short-lived species, the rapid takeover of mutants once they appear, and the emergence of single dominant deletions per cell, thereby resolving many inconsistencies of previous models.

The accumulation of deletion mutants has important physiological consequences. In skeletal muscle, clonal expansions underlie the segmental cytochrome oxidase-deficient regions known as ragged-red fibres. Studies in humans, rhesus monkeys, and rodents have shown that once mutant loads exceed 80–90%, affected fibres display marked atrophy, splitting, and eventually fibre loss, directly contributing to sarcopenia^2,5,12-15^. In neurons of the substantia nigra, clonal deletions are abundant in aged individuals and Parkinson disease patients, with deletion burdens exceeding 60% tightly associated with COX deficiency^1,3,6^. These findings highlight not only the causal role of mtDNA deletions in age-related tissue decline but also their potential contribution to neurodegeneration.

The temporal dynamics of clonal expansion are therefore of central interest. Does the accumulation of mutants proceed slowly and stochastically throughout life, or do specific mechanisms trigger a rapid takeover once deletions arise in vulnerable genomic regions? Understanding these dynamics is critical, as they determine how long cells can tolerate mutant genomes before crossing the pathogenic threshold. The transcription-coupled replication model predicts a biphasic process: an initial lag until a deletion removing the feedback gene arises, followed by a rapid exponential expansion phase. This would result in relatively similar accumulation times across mammalian species despite differences in lifespan^27^.

In this study, we propose a novel approach to infer accumulation times of mtDNA deletion mutants from cross-sectional single-cell RNA sequencing (scRNAseq) data. Although our primary interest lies in DNA-level deletions, we use transcriptomic data as a proxy because comprehensive single-cell mtDNA sequencing data are currently unavailable at sufficient scale and accuracy. Since deletions in the mitochondrial genome directly eliminate the corresponding transcripts, patterns in mitochondrial gene expression provide an indirect but informative readout of the underlying mutational landscape. By analysing these patterns across large numbers of cells, it becomes possible to extract quantitative information about mutant accumulation dynamics that cannot be accessed by longitudinal measurements.

The Results section is organised as follows. In the first section, ‘*Obtaining longitudinal from cross-sectional scRNAseq data’*, we introduce the conceptual framework that links single-cell transcriptomic measurements to underlying mutant accumulation trajectories and illustrate how cross-sectional snapshots can encode information about longitudinal dynamics. In the next section, ‘*Generation of synthetic data’*, we describe two stochastic models of the mitochondrial life cycle that incorporate replication, mutation, and degradation, and use these models to generate synthetic datasets that serve as gold standards for method validation. The advantage of using synthetic data is that the true parameters underlying the system are known, allowing us to rigorously test the accuracy of inference methods. We then analyse an additional key observable in the section ‘*Distribution of unique mutants’*, where we investigate how many distinct deletion species are expected to coexist within individual cells. Using the synthetic data, we show that the number of unique mutant types follows a Poisson distribution and examine how its single parameter depends on mutation rate, selection advantage, and age. In the subsequent section, ‘*Using the Moran process to model accumulation times’*, we demonstrate that a reduced stochastic framework based on the Moran process^28-30^ accurately reproduces the accumulation time distributions obtained from the more detailed simulations. Finally, in ‘*Fitting the Moran process to real world data’*, we show how the Moran model can be fitted to cross-sectional summary statistics, enabling the extraction of mutation probabilities, selection advantages, and associated confidence intervals in a way that is directly applicable to experimental scRNAseq data.

Together, these results establish a coherent pipeline that links single-cell transcriptomic measurements to quantitative estimates of mtDNA mutant dynamics, providing the foundation for the experimental application presented in a companion study.

## Results

### Obtaining longitudinal from cross-sectional scRNAseq data

Measuring the amount of mitochondrial DNA (mtDNA) mutants inside a single cell inevitably destroys that cell, making it impossible to obtain accumulation times by repeatedly following the same cell over time. This raises the critical methodological question of whether it is possible to derive such information from cross-sectional data, such as single-cell RNA sequencing (scRNAseq). Ideally, one would like to obtain direct quantitative measurements of mutant and wild-type genomes at the DNA level. Single-cell DNA sequencing (scDNAseq) would in principle provide the necessary resolution, and indeed, numerous areas of biology, from cancer genomics to developmental lineage tracing, would benefit from such approaches^31^. However, technical obstacles have historically limited the reliability of scDNAseq. Problems such as incomplete genome coverage, amplification bias, allelic dropout, and high error rates have restricted its widespread use, especially in applications where quantitative accuracy is essential. Recent methodological innovations have begun to address these challenges, for instance through intracellular genomic amplification strategies that amplify DNA within intact, permeabilized cells, thereby avoiding some of the bottlenecks of traditional single-cell DNA protocols^32^. Despite these promising advances, scDNAseq is still not yet available at the scale and accuracy required to systematically investigate mtDNA deletions across large numbers of individual cells.

Given these limitations, single-cell transcriptomics provides a practical and informative alternative. The rationale is that deletions at the DNA level are faithfully reflected at the RNA level. This creates a direct link between DNA-level deletions and their transcriptional consequences. Thus, the relative abundance of mutant versus wild-type mtDNA can be approximated by measuring the ratio of transcripts that contain a deletion versus those that don’t. While not a perfect proxy, since RNA stability and processing may introduce additional layers of complexity, this approach nonetheless offers a unique window into the intracellular dynamics of mtDNA populations. Importantly, scRNAseq can be applied at high throughput, providing data from tens of thousands of cells in parallel, which is essential to capture the rare and stochastic events underlying clonal expansion of mtDNA deletions.

Fig. 1 illustrates the conceptual framework using three example cells, plotted with time on the horizontal axis. Initially, all cells contain exclusively wild-type mtDNA molecules. At each round of replication, however, there is a finite probability that a deletion mutation occurs. This generates the first mutant molecule, which can then expand within the cell population of mtDNAs. Over time, the mutant genomes may clonally dominate, progressively displacing the wild type. This process is symbolized in the inset diagram, with wild-type genomes shown in red and mutant genomes in green, loosely following simulation results from^26^. If a scRNAseq experiment is performed at a given time point (marked by the vertical blue bar), the mtDNA composition of each cell is captured indirectly through its transcriptome. In the example shown, cell 1 has undergone complete takeover by mutant genomes and therefore produces only mutant-associated transcripts. Cell 2 has not yet experienced a mutation event and thus shows only wild-type signals. Cell 3 represents an intermediate state, with transcripts derived from both wild-type and mutant genomes, reflecting a cell in the midst of the accumulation process.

**Fig. 1.**
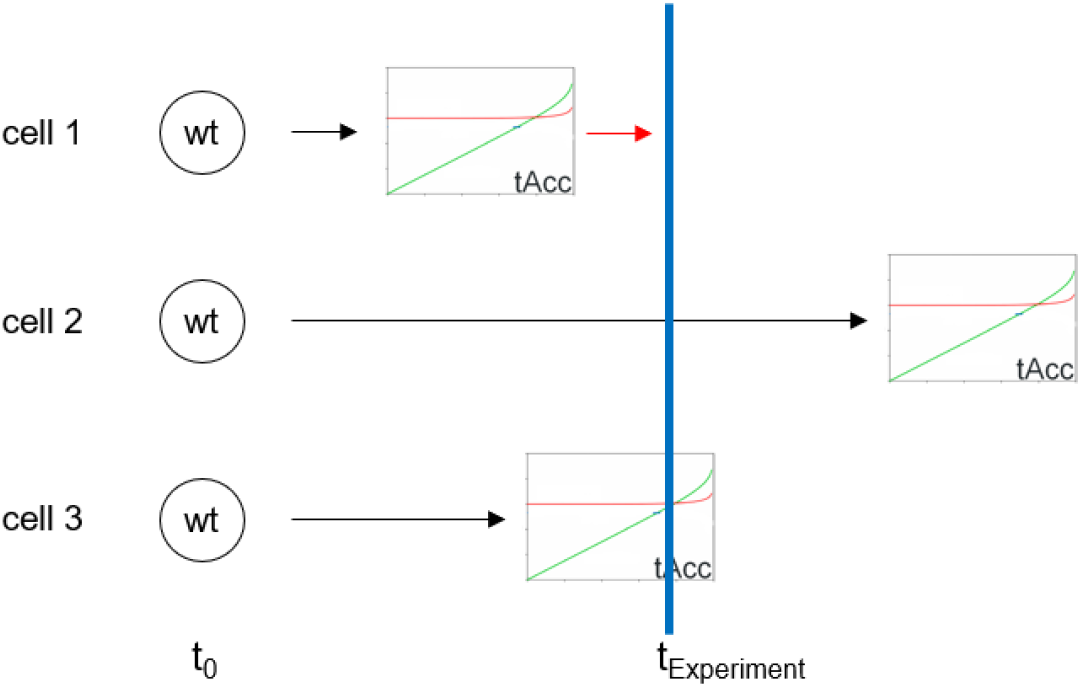
Individual cells start with only wild type mtDNA, but there is a certain probability per replication, *mutProb*, that a mutation occurs. From that moment on mutants (green) accumulate until the cell is taken over after tAcc days. tAcc can be a fixed number or come from a distribution. At time tExp (blue bar), a single cell NGS experiment is performed, which in this case would reveal that cell1 is completely taken over (but did not undergo apoptosis), cell2 is still wild-type and cell3 is caught in the middle of mutant accumulation.

This schematic makes clear that the probability of observing cells at different stages of mutant takeover depends on a set of underlying parameters, such as the mtDNA half-life, the mutation rate, and any selective advantage carried by the deletion mutants. Crucially, this also implies that, given sufficient single-cell transcriptomic data, one can invert the problem and deduce the relevant parameters from the distribution of mutant-to-wild-type ratios observed across many cells. In this way, scRNAseq data provide not only a snapshot of the heterogeneity of mtDNA populations across individual cells but also a means to extract quantitative information about the fundamental processes governing the accumulation of mitochondrial mutants.

### Generation of synthetic data

To test whether accumulation times of mtDNA mutants can indeed be inferred from cross-sectional data, we simulated the mitochondrial life cycle, comprising replication, mutation, and degradation, using two distinct stochastic models. This strategy provides access to longitudinal trajectories of mutant expansion, which are experimentally inaccessible. From the simulated data we obtained distributions of mutant accumulation times that serve as a gold standard against which our analytical approach can be evaluated.

Model-1, originally developed in^26^, consists of three ordinary differential equations (ODEs) describing the time evolution of wild-type (wt) and mutant (mt) mtDNAs as well as ATP. In this formulation, the cell strives to maintain a fixed ATP level, which results in an expanding mtDNA population size as mutant genomes accumulate. Model-2 represents a streamlined variant of Model-1 and involves only two equations, one each for wild-type and mutant mtDNAs. Here the regulation is imposed at the level of total mtDNA copy number, i.e. the cell maintains a constant population size, N, throughout the accumulation process, independent of ATP dynamics.

In both models mutant accumulation arises from the assumption that mutant genomes replicate at a higher rate than wild-type (*selAdv* > 0). A detailed discussion of the biological justification for these assumptions, as well as the parameter values used (e.g. N=1000, mtDNA half-life=10 days), is provided elsewhere^26,27^. Although the ODE formulation is mathematically concise, we implemented the models as stochastic simulations in Java to capture random mutation events and probabilistic replication and degradation. The simulation code is available from our GitLab repository. This stochastic framework also allows us to account for different classes of deletion mutants. Based on our earlier predictions, only those deletions that disrupt genes involved in the proposed transcriptional feedback mechanism confer a selective replication advantage, while deletions in other regions provide no advantage and can only persist by neutral drift^26^. Consequently, the program explicitly tracks wild-type genomes, mutants with a replication advantage, and mutants without advantage.

**Model-1**

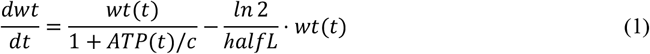

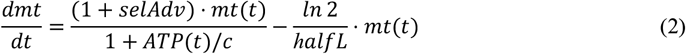

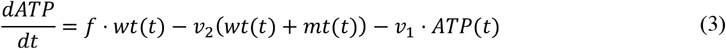

**Model-2**

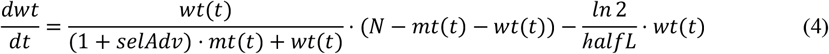

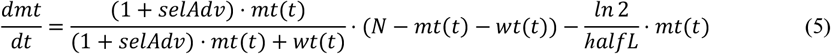

Employing two models with distinct regulatory assumptions allowed us to assess the robustness of our analytical method. Since the true mechanisms controlling mitochondrial biogenesis and turnover remain incompletely understood, it is important to verify that the inference procedure is not overly sensitive to model-specific details. For both models we simulated the fate of 100000 cells over a three-year period, approximating the lifespan of a mouse, and recorded the time from the first appearance of a mutant until takeover by mutants. Simulations varied the selection advantage (*selAdv*), the mutation probability per replication (*mutProb*), and the fraction of mutations generating an advantageous mutant (*fracAdv*). Takeover was defined as the point at which mutants reached 90% abundance; in Model-1 this coincided with ATP collapse (ATP = 0), while in Model-2 it was explicitly imposed as the 90% threshold.

Fig. 2 summarizes the resulting distributions of accumulation times for different parameter settings. Please note that these times start from the occurrence of the first mutant mtDNA molecule and not from birth of the animal. Histograms are binned in 30-day intervals, corresponding to the snapshot frequency of the simulation. For cases with *fracAdv* = 0.2 (Fig. 2B,D), we increased the mutation rate from 6.8e-6 to 3e-5 per replication to ensure sufficient numbers of takeover events among the 100000 simulated cells; otherwise, the number of affected cells would have been too low to yield reliable histograms.

**Fig. 2.**
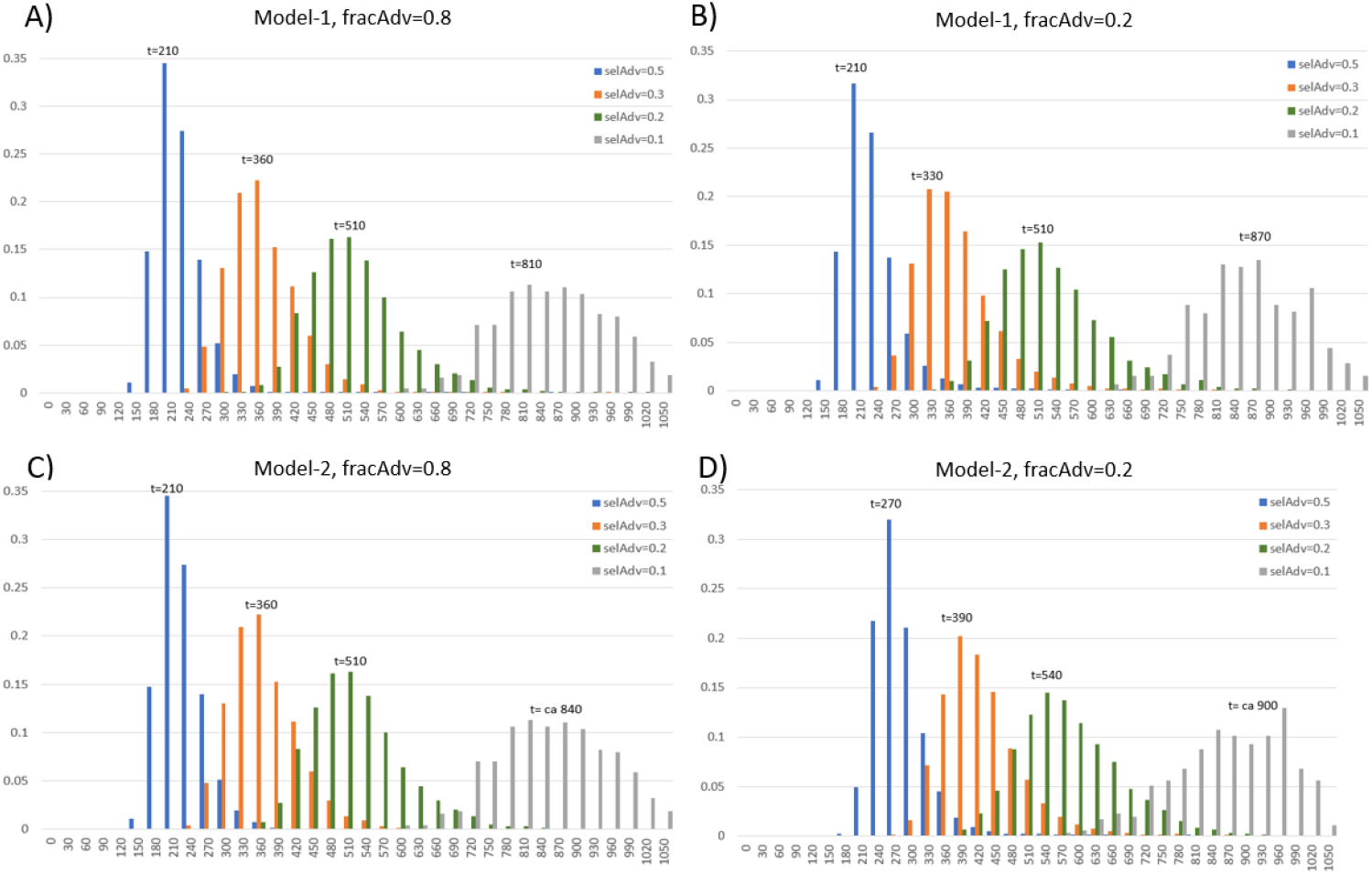
Distributions of accumulation times for mitochondrial deletion mutants (from first occurrence until 90% prevalence) obtained from stochastic simulations. Panels A–D show results from two different simulation models of the mitochondrial life cycle (see main text). Each histogram represents the distribution of accumulation times. Panels A and C depict simulations where 80% of mutations confer a selective advantage, while Panels B and D show simulations where only 20% of mutations produce advantageous mutants.

Overall, the results show that accumulation times are approximately normally distributed with a long right-hand tail. As expected, higher selection advantages shortened the takeover time. Notably, the two models produced very similar distributions (compare Fig. 2A,B with Fig. 2C,D), suggesting that the precise details of mitochondrial regulation, whether based on ATP homeostasis or fixed copy number, are not critical determinants of mutant accumulation kinetics. Likewise, varying the fraction of mutants with selective advantage (*fracAdv*) had little impact on the distributions. In fact, inspection of the simulation data revealed that none of the takeover events was caused by mutants without selective advantage; these genomes never succeeded in displacing wild-type mtDNAs. This outcome is consistent with our earlier finding that accumulation via drift alone is not feasible in short-lived species such as mice or rats^25^.

### Distribution of unique mutants

Using the synthetic datasets, we can not only extract accumulation times for deletion mutants, as shown in the previous section, but also characterize an equally important feature of mitochondrial mutational dynamics: the distribution of unique mutant species present in individual cells at old age. As noted earlier, post-mitotic cells typically harbour only one, or at most a very small number, of different deletion types. This empirical observation is a major constraint on any mechanistic hypothesis seeking to explain mtDNA mutant accumulation, because several proposed models, most notably neutral drift, predict a much larger diversity of mutants within single cells, especially in short-lived species^25,26^. Understanding why only a few deletion species dominate is therefore essential for distinguishing between competing hypotheses.

To explore this, we analysed the state of each simulated cell at the end of a 3-year simulation period (corresponding to the lifespan of a mouse) across a range of parameter combinations for selection advantage (*selAdv*) and the fraction of mutations that lead to advantageous mutants (*fracAdv*). For each parameter set, we counted how many distinct mutant types were present in every cell. The resulting distributions, shown in Fig. 3, reproduce earlier observations^26^, i.e. whenever deletion mutants possess a selection advantage, the overwhelming majority of cells contain only a very small number of unique deletion species.

**Fig. 3.**
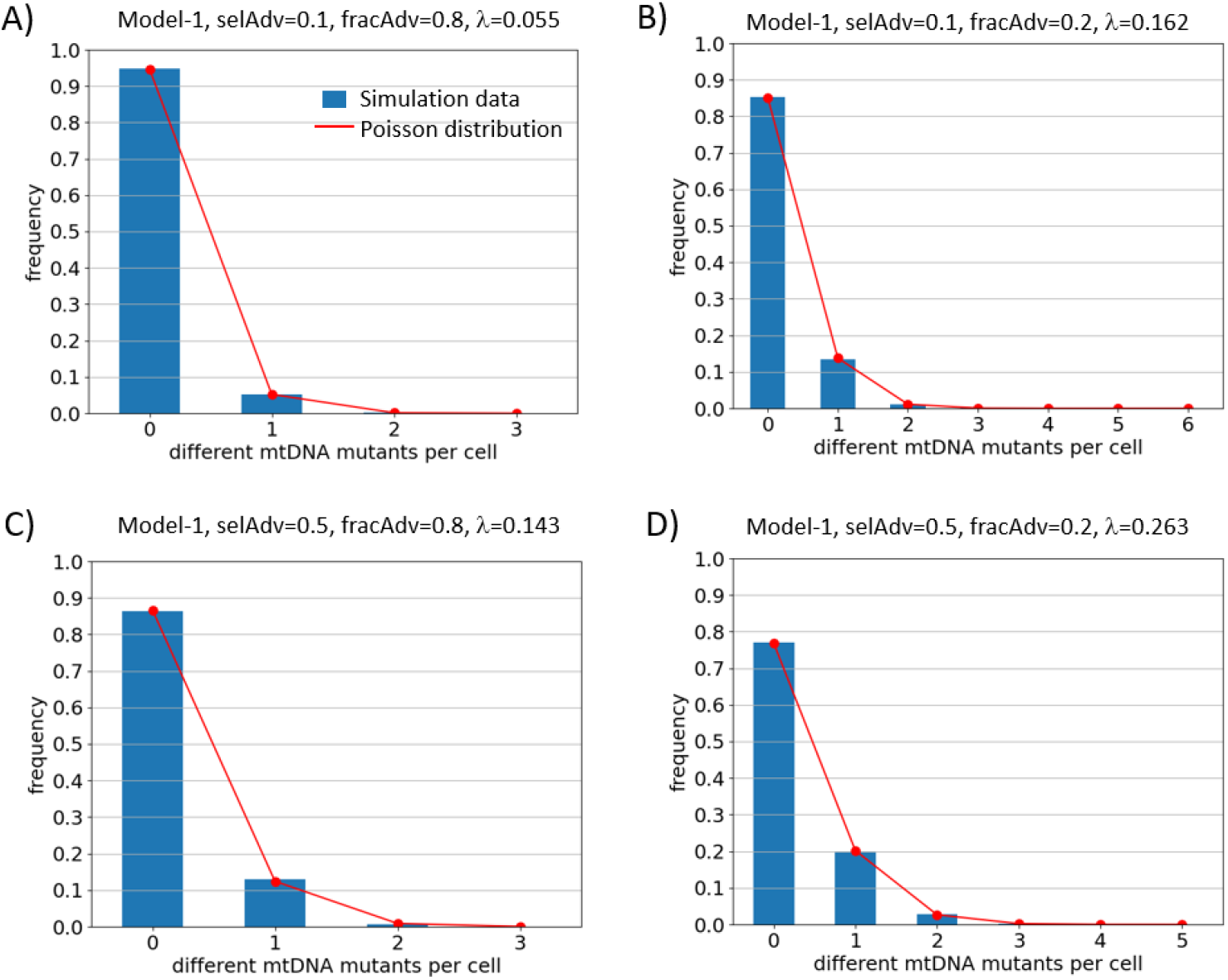
Distributions of the number of unique deletion mutants per cell at the end of a three-year mouse lifespan, obtained from stochastic simulations of Model-1. A value of zero indicates cells containing only wild-type mtDNAs. Each panel corresponds to a different combination of selection advantage (*selAdv*) and fraction of advantageous mutants (*fracAdv*). Blue bars show the simulation results, while red lines indicate the corresponding Poisson distributions calculated using the mean number of unique mutants observed in each simulation.

Because the appearance of deletion mutants is driven by rare mutation events occurring during replication, we next asked whether the number of unique deletion types per cell might follow a Poisson distribution. The Poisson process is the canonical model for rare, independent events occurring at a constant rate, and it is therefore a natural candidate for describing the stochastic generation of deletion species. To test this idea, we computed the mean number of unique mutants across the 100000 simulated cells for each parameter condition, and used this mean as the Poisson rate parameter (λ). We then overlaid the resulting Poisson probability mass function (in red) onto the histogram of observed values (blue bars) in Fig. 3. The agreement is remarkably close across all parameter sets examined. At first glance, it may appear intuitive that rare mutation events would produce a Poisson-distributed number of unique deletion types. However, the excellent agreement observed here is far from obvious. The mitochondrial life cycle is not a simple accumulation of independent mutation events, instead it includes several nonlinear processes that could in principle distort or completely break the Poisson pattern. First, once a deletion mutant arises, it undergoes replication, competition, and degradation within the mitochondrial population. Mutants with a selective advantage tend to expand rapidly, effectively suppressing the fixation of other mutants arising later. This introduces interactions between mutant lineages, violating the independence assumption of a pure Poisson process. Second, degradation (mitophagy) and the feedback-regulated replication dynamics of wild-type and mutant genomes further influence the fate of both mutant classes and their coexistence. These processes introduce time-dependent biases, making it unclear whether the number of unique mutants should retain a simple distributional form. Third, the presence of two classes of mutants, with and without selection advantage, adds additional structure, as the former tend to dominate the population while the latter persist only transiently.

A direct consequence of the finding that the distribution of unique mutant types follows a Poisson distribution is that it can be fully characterized by a single parameter, λ, representing the mean number of distinct deletion species present per cell. This greatly simplifies the description of mutational diversity and reduces the problem to understanding how λ depends on the underlying biological parameters of the system. To investigate this, we systematically varied the key model parameter, selection advantage (*selAdv*), fraction of advantageous mutations (*fracAdv*), mutation probability (*mutProb*), and age, and computed λ for each condition based on the synthetic data. The resulting relationships are shown in Fig. 4.

**Fig. 4.**
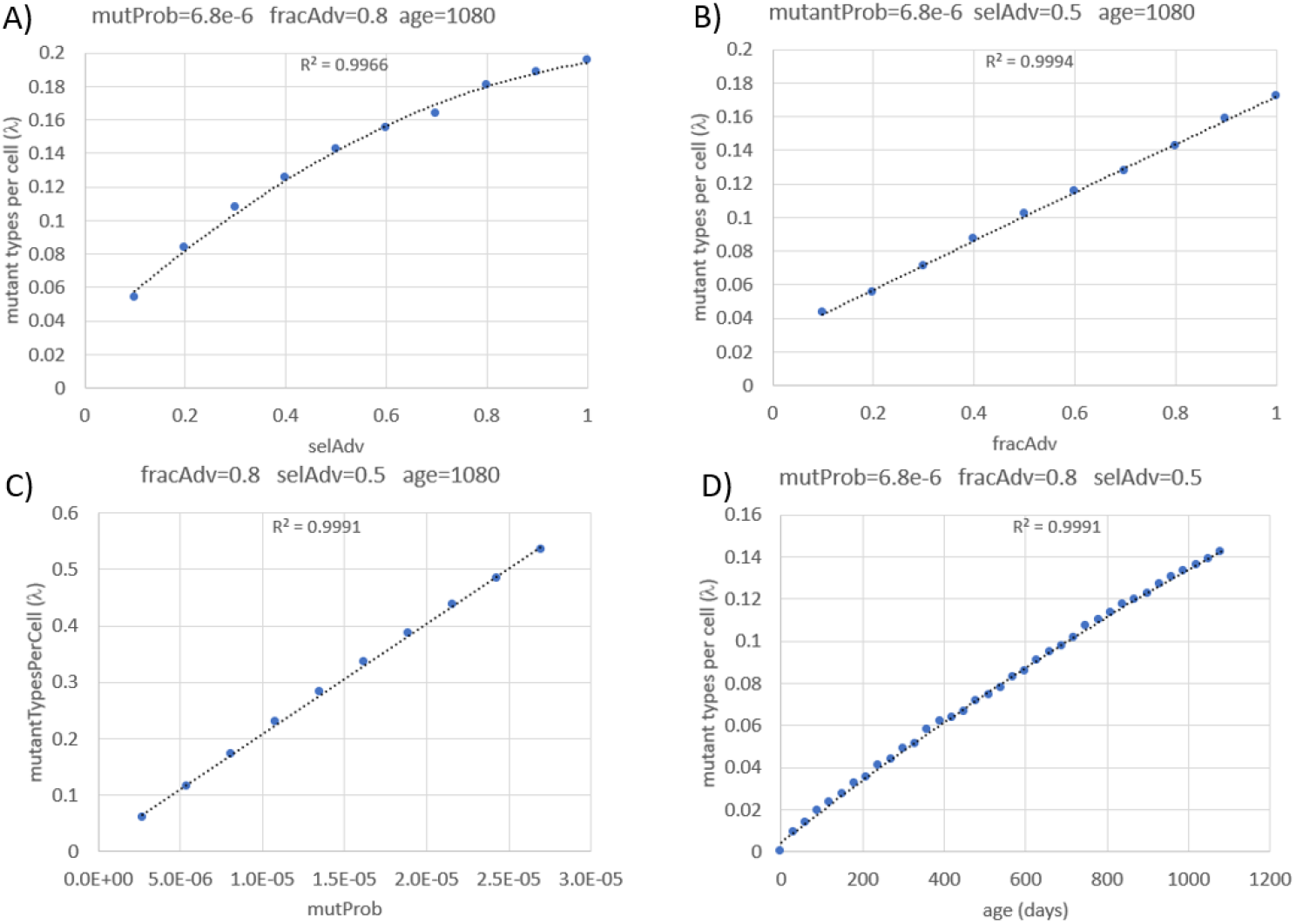
Dependence of the mean number of unique deletion mutants (λ) on model parameters. Linear fits highlight the nearly perfect proportional dependence of λ on *fracAdv* and *mutProb*, whereas quadratic trendlines capture the nonlinear relationships with *selAdv* and age.

The mean number of unique mutants λ increases almost perfectly linearly with both the fraction of advantageous mutants (*fracAdv*) and the mutation probability (*mutProb*). In contrast, λ exhibits a nonlinear, nearly quadratic dependence on both selection advantage (*selAdv*) and age. These relationships suggest that the number of unique deletion species in a cell results from a balance between stochastic mutational input (linear effects) and the amplification dynamics driven by selection and time (non-linear effects). The observation that such clear mathematical patterns emerge from a complex biological model underscores the robustness of the underlying processes and suggests that similar scaling laws may hold *in vivo*.

### Using the Moran process to model accumulation times

The synthetic data generated in the previous section provide a test-case for real scRNAseq measurements, allowing us to test whether it is possible to recover key parameters of the model. Because the true parameter values are known in this case, the synthetic data serve as a gold standard for evaluating the accuracy of our inference approach.

To perform this inference we used the Moran process^28-30^, a classical stochastic model from population genetics that that describes how a new variant spreads in a finite population of constant size. The mathematical details of the model, including the derivation of accumulation time distributions using recursive first-step analysis, are provided in the Methods section. For comparison with the synthetic data we computed the distribution of accumulation times under the Moran process. Instead of requiring full fixation, we stopped the calculation once mutants reached 90% of the population, which corresponds to our operational definition of takeover in the simulations.

The distributions derived from the Moran model (Fig. 5) closely match those obtained from the stochastic simulation models (Fig. 2). Only in the case of a very small selection advantage (*selAdv* = 0.1) do noticeable discrepancies appear, i.e. the synthetic data show a peak probability shifted toward earlier times. This difference is not due to a failure of the Moran model but rather to an observational bias in the simulations. Because the simulations were limited to three years, the approximate lifespan of a mouse, extremely long takeover times could not be captured. Truncating the distribution in this way artificially inflates the probabilities of earlier takeover. In contrast, the Moran-based calculation was extended until the entire distribution was observed.

**Fig. 5.**
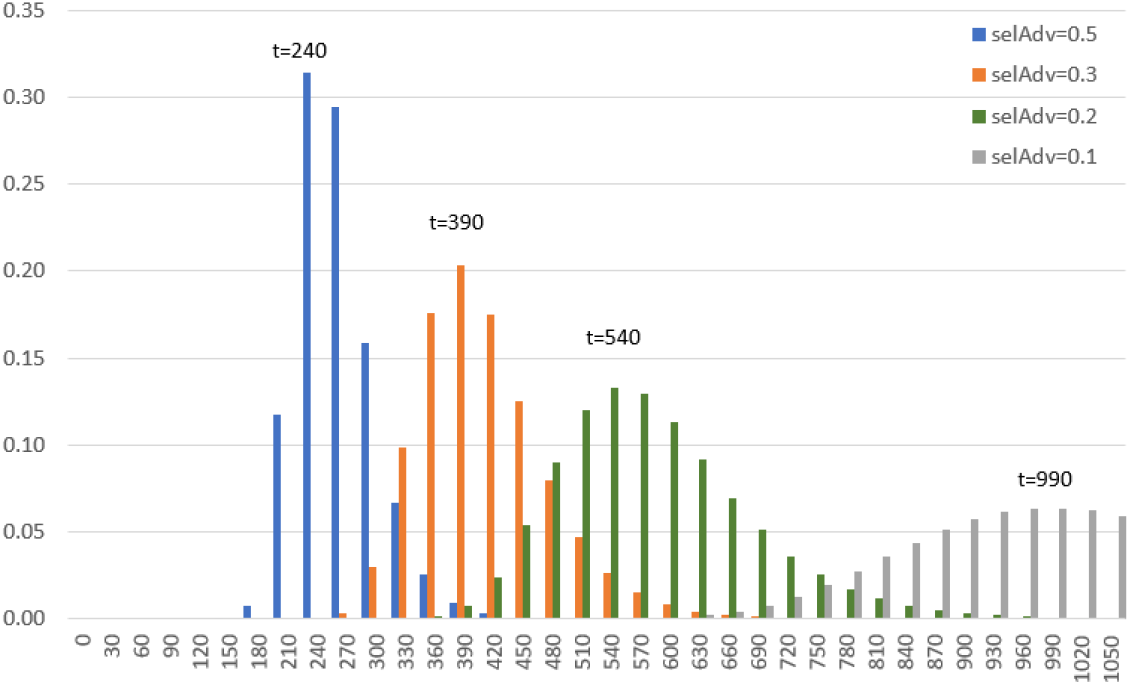
Distributions of accumulation times for mitochondrial deletion mutants (from first occurrence until 90% prevalence) based on the Moran model. Each histogram represents the distribution of accumulation times for a certain selection advantage.

These results demonstrate that the Moran process can successfully reproduce the accumulation time distributions generated by two different, more detailed, stochastic simulations. Despite the fact that the true regulation of mitochondrial biogenesis and turnover is not fully understood, this agreement suggests that the Moran model provides a robust and tractable framework for analysing the accumulation dynamics of mitochondrial mutants. Importantly, while synthetic data allow direct measurement of accumulation times by following individual simulated cells, this is not possible experimentally because cells are destroyed during scRNAseq measurement. In the next section we therefore show how the Moran model can be exploited to infer the selective advantage of mtDNA mutants from cross-sectional data, thereby bridging the gap between theory and experiment.

### Fitting the Moran process to real world data

Longitudinal single-cell trajectories, such as those obtained in the previous section from simulations, are not available in experimental scRNAseq data. Instead, we only have cross-sectional information so that we can measure the fraction of cells that harbour different proportions of deletion mutants (or, equivalently, transcripts with deletions) only at specific time points. This information can also be derived from our synthetic data as well as the Moran process (see Fig. 6). To demonstrate this, we utilized the forward master equation of the Moran process to compute the probability distribution over mutant counts at any time *t*. As detailed in the Methods section, this approach captures continuous turnover and competition among genome classes, mapped to physical time units. From this, summary measures can be extracted such as the fraction of cells with more than a given percentage of mutants.

**Fig. 6.**
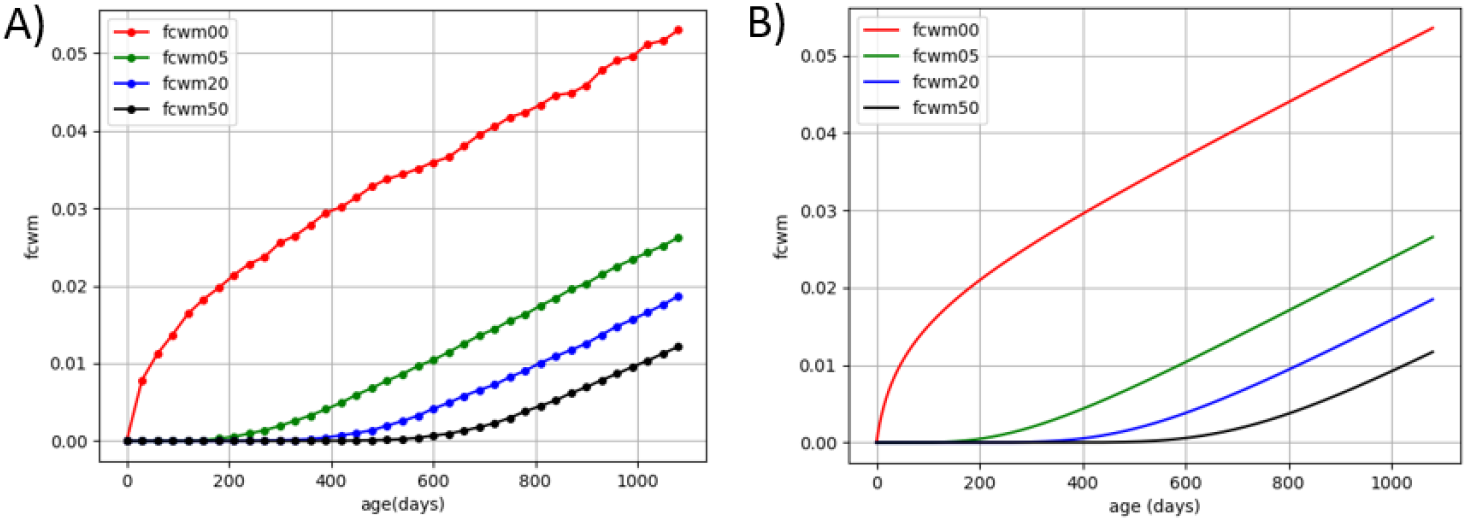
A) Fraction of cells with more than 0%, 5%, 20% and 50% of mutant mtDNAs as function of time for Model-1 using the parameter values *selAdv*=0.1, *mutProb*=6.8e-6 and *fracAdv*=0.8. Data values are exported and displayed in 30 day intervals. B) The same type of information generated from the forward-master equation of the Moran process.

Fig. 6B shows that when the true parameter values are used, the Moran process reproduces the synthetic data curves from Fig. 6A with high accuracy. For real experimental data, however, the parameter values are not known a priori and must instead be inferred by fitting the model to the observed data. We therefore performed parameter estimation by fitting three key parameters, mutation probability (*mutProb*), selection advantage (*selAdv*), and the fraction of advantageous mutants (*fracAdv*), to the data. This was achieved by minimizing the sum of squared differences between the Moran predictions and the empirical curves using a differential evolution algorithm (see Methods).

Table 1 summarizes the ground truth parameter values together with the estimated mean, coefficient of variation (CV), and 95 % confidence intervals obtained via parametric bootstrapping. For all three parameters, the estimated means are very close to the true values used to generate the data, indicating that the fitting procedure is unbiased in this parameter regime. The CV values are small, in particular for the mutation probability *mutProb* and the selection advantage *selAdv*, demonstrating high statistical precision.

**Table 1:** Parameter estimates obtained via parametric bootstrapping (N=40) of 100000 simulated cells.

| Name | Ground Truth | Estimate Mean | Estimate CV (%) | 95% CI |
| --- | --- | --- | --- | --- |
| *mutProb* | 6.8e-6 | 6.74e-6 | 1.71 | [6.55e-6, 6.99e-6] |
| *selAdv* | 0.1 | 0.108 | 4.51 | [0.094, 0.117] |
| *fracAdv* | 0.8 | 0.87 | 7.41 | [0.71, 1] |

The fraction of advantageous mutations (*fracAdv*) shows a larger CV and an asymmetric confidence interval, with the upper bound reaching the imposed maximum value of 1. This behaviour indicates partial non-identifiability of *fracAdv* near its upper boundary. Once the fitting procedure reached a large value for *fracAdv*, the data contain limited information to distinguish, for example, *fracAdv* = 0.9 from *fracAdv* = 1. Importantly, this does not reflect a failure of the fitting procedure, but rather a structural limitation of the available observables under this parameter regime.

Fig. 7 shows histograms of the estimated parameter values obtained from the 40 bootstrap data sets (with *mutProb* displayed on a logarithmic scale). All three distributions are unimodal and approximately symmetric around a single maximum, with no evidence for secondary optima or multimodality. This indicates that the optimization landscape is well behaved and that the stochastic optimization procedure converges consistently to the same region of parameter space. The relatively narrow spread of the distributions further confirms the high precision of the estimates for data sets of this size.

**Fig. 7.**
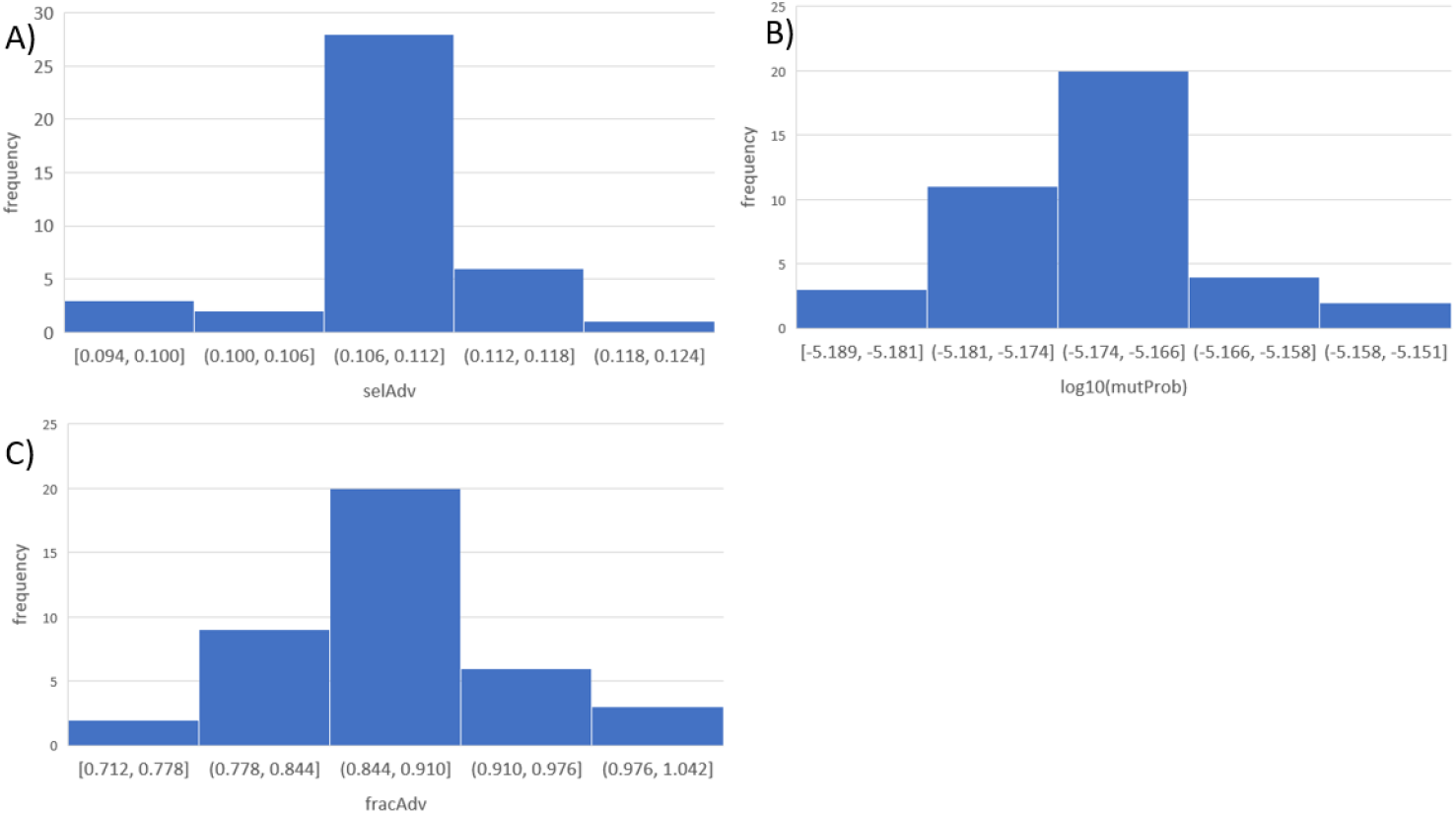
Histograms show the estimated values of A) selection advantage (*selAdv*), B) mutation probability (*mutProb*), and C) fraction of advantageous mutations (*fracAdv*) obtained from 40 independent bootstrap data sets, each based on 100000 simulated cells. The mutation probability is displayed on a logarithmic scale. All distributions are unimodal and centred close to the corresponding ground truth values, indicating unbiased estimation and stable convergence of the fitting procedure.

In an forthcoming publication^33^, we apply the same fitting framework to experimental single-cell transcriptomics data, which are typically based on substantially smaller numbers of cells, often in the range of a few thousand. To assess how parameter uncertainty scales with reduced sample size, and to obtain confidence interval estimates that are relevant for such experimental settings, we repeated the parametric bootstrapping procedure using synthetic data sets based on only 10000 simulated cells.

The results are summarized in Table 2. As expected, the estimated means remain close to the ground truth values, demonstrating that the fitting procedure remains unbiased even at lower cell numbers. However, the CV values increase and the confidence intervals widen for all three parameters, reflecting the higher stochastic variability inherent to smaller sample sizes. The key idea underlying this analysis is that, when fitting real experimental data with a comparable number of cells, the confidence intervals obtained from synthetic data of the same size provide a quantitative estimate of the expected statistical uncertainty. Since the synthetic data are generated from the same stochastic model that underlies the inference procedure, these confidence intervals can be interpreted as model-conditional minimal uncertainty estimates for the experimental parameter fits. Of course, biological and experimental variability can increase these estimates further.

**Table 2:** Parameter estimates obtained via parametric bootstrapping (N=40) of 10000 simulated cells.

| Name | Ground Truth | Estimate Mean | Estimate CV (%) | 95% CI |
| --- | --- | --- | --- | --- |
| *mutProb* | 6.8e-6 | 6.65e-6 | 4.24 | [6.11e-6, 7.22e-6] |
| *selAdv* | 0.1 | 0.107 | 9.74 | [0.089, 0.125] |
| *fracAdv* | 0.8 | 0.87 | 13.03 | [0.65, 1] |

Finally, to demonstrate the robustness of the fitting approach across different regions of parameter space, we repeated the entire procedure (again based on 10000 simulated cells) using a second set of parameter values that differs substantially from the first (higher mutation probability, stronger selection, and a lower fraction of advantageous mutations). The results are shown in Table 3. As in the previous analyses, the estimated means agree well with the corresponding ground truth values, indicating that the fitting procedure generalizes well across parameter regimes. Notably, the confidence interval for *fracAdv* is considerably narrower in this case. This can be explained by the fact that *fracAdv* = 0.2 lies well within the interior of the admissible parameter range, far from the upper boundary at 1. Consequently, the data contain sufficient information to constrain *fracAdv* more tightly, and the identifiability issues observed in the high-*fracAdv* regime are no longer present. Another noteworthy observation is that the upper confidence bound for *mutProb* lies slightly below the true value. This behaviour is consistent with finite-sample variability in the bootstrap procedure and reflects a small downward bias of the estimator under this specific parameter combination. Importantly, the true value still lies close to the confidence interval, and repeated bootstrap analyses with different random seeds yield similar results, indicating that this effect is modest and does not compromise the overall robustness of the inference.

**Table 3:** Parameter estimates obtained via parametric bootstrapping (N=40) of 10000 simulated cells for a 2nd set of parameter values.

| Name | Ground Truth | Estimate Mean | Estimate CV (%) | 95% CI |
| --- | --- | --- | --- | --- |
| *mutProb* | 3e-5 | 2.9e-5 | 1.78 | [2.80e-5, 2.99e-5] |
| *selAdv* | 0.5 | 0.538 | 8.05 | [0.461, 0.628] |
| *fracAdv* | 0.2 | 0.21 | 8.14 | [0.18, 0.24] |

In summary, these results demonstrate that by fitting the Moran model to cross-sectional data, as would be available from scRNAseq experiments, it is possible to recover parameters that provide important insights into the occurrence and accumulation of mtDNA deletion mutants. The estimates of selection advantage in particular provide direct insight into whether mutant accumulation is driven by random drift or positive selection (which allows to differentiate between different hypotheses), and they can be used to calculate the expected distribution of accumulation times from the first appearance of a mutant until its eventual takeover of the mitochondrial population.

## Discussion

The accumulation of mitochondrial DNA (mtDNA) deletion mutants in post-mitotic cells has long been recognized as a hallmark of ageing^1-11^. Numerous studies in skeletal muscle, neurons, and other tissues across mammals have shown that mtDNA deletions, although rare in early life, can clonally expand within single cells to levels that surpass the threshold required for respiratory chain dysfunction. Once mutant loads exceed approximately 80–90%, oxidative phosphorylation collapses, leading to cytochrome c oxidase (COX) deficiency, atrophy, and ultimately the loss of affected cells. These events are central to age-related sarcopenia and neurodegeneration. Yet, despite decades of research, the fundamental question of why certain deletion mutants are able to outcompete wild-type genomes and accumulate to such high levels has remained unresolved.

Several mechanistic hypotheses have been advanced. Early “vicious cycle” models proposed that respiratory dysfunction increased reactive oxygen species (ROS) production, thereby generating further mtDNA damage^20,21^. While appealing, this idea is inconsistent with the observation that single cells typically harbour a dominant deletion species rather than a heterogeneous mixture of multiple mutants. Other models emphasized structural or functional advantages of mutants. The “survival of the smallest” hypothesis suggested that shorter genomes replicate more rapidly, whereas the “survival of the slowest” model argued that dysfunctional mitochondria escape turnover by producing fewer damaging ROS. However, both models face serious conceptual and empirical limitations. Genome size differences appear too small to meaningfully affect replication kinetics, and mitophagy mechanisms generally target rather than protect dysfunctional organelles.

Neutral drift offered a more parsimonious explanation. Mathematical models demonstrated that random replication and degradation under relaxed control could, in principle, generate clonal expansions of mtDNA mutations^16,17^. Drift is particularly effective when copy numbers are low and sufficient time is available for stochastic fluctuations to carry a mutant to dominance. These models successfully reproduced the frequency of COX-deficient cells in long-lived species such as humans. However, they fail for short-lived species like mice or rats, where drift would predict multiple mutant species per cell, a pattern not observed experimentally. This discrepancy highlighted the need for an additional selective mechanism.

To address this gap, a transcription-coupled replication model was proposed^26,27^. In mammalian mitochondria, replication is primed by transcripts, and transcription is proposed to be subject to negative feedback once sufficient respiratory chain subunits are produced. Deletions that remove the genes responsible for this feedback loop escape regulation, leading to persistently high transcription and replication initiation. Mutants encompassing the ND4 or ND5 genes, repeatedly identified in expanded deletions across species, are therefore predicted to gain a cis-acting replication advantage. Computational models incorporating this mechanism explain both the low heteroplasmy levels observed in short-lived animals and the dominance of single deletions per cell, while still remaining consistent with patterns in long-lived species.

The key prediction of this framework is that once such a mutant arises, its accumulation time from first occurrence until cellular takeover should be relatively short, on the order of several months, and similar across species. Knowing this accumulation time is therefore crucial for distinguishing between competing hypotheses of mutant expansion. Unfortunately, the accumulation time cannot be measured directly, since experimental approaches to quantify mtDNA mutants destroy the cell. Longitudinal measurements on the same cell are therefore impossible. In this study, we addressed this challenge by proposing that accumulation times can be inferred indirectly from cross-sectional measurements such as single-cell RNA sequencing. Although scRNAseq captures mRNA rather than DNA, deletions in the genome are faithfully reflected in the transcriptome. Thus, the ratio of mutant to wild-type transcripts provides a proxy for the underlying genome composition. With sufficient sampling across many cells, it should be possible to reconstruct the distribution of accumulation states and, from this, infer the parameters governing the expansion process.

Using two stochastic models of the mitochondrial life cycle, we generated synthetic datasets that serve as a gold standard. Despite differing assumptions about mitochondrial regulation, ATP homeostasis versus fixed copy number, both models produced nearly identical distributions of mutant accumulation times. This demonstrates that the detailed regulation of mitochondrial turnover is not critical for the macroscopic dynamics of mutant expansion. Crucially, only mutants with a selective replication advantage were able to achieve takeover, reinforcing the conclusion that drift alone is insufficient in short-lived species.

Beyond accumulation times, we analysed the distribution of unique mutant species per cell, a highly informative observable. Experimentally, post-mitotic cells are typically dominated by one or very few deletion species. Our simulations reproduced this pattern whenever a subset of mutants possessed a selection advantage. Remarkably, the number of unique mutants per cell followed an almost perfect Poisson distribution, characterized by a single parameter λ. This finding is nontrivial since the mitochondrial life cycle includes selection, competition, and degradation processes that could in principle distort such a simple statistical structure. The persistence of a Poisson distribution indicates that the stochastic generation of rare mutation events is the dominant determinant of mutant diversity, while selection primarily filters which of these mutants persist.

Systematic variation of model parameters revealed that λ depends linearly on the mutation probability and the fraction of advantageous mutations, but shows a nonlinear, approximately quadratic dependence on selection advantage and age. These scaling relationships provide additional mechanistic insight, suggesting that mutational input governs diversity in a proportional manner, whereas selection and time amplify early-arising mutants and suppress later competitors in a nonlinear fashion. The emergence of such simple mathematical relationships from a biologically complex system underscores the robustness of the underlying processes and suggests that similar patterns may be observable in real data.

To connect these insights to experimental measurements, we employed the Moran process as a tractable mathematical framework for modelling mutant dynamics^28-30^. The Moran model accurately reproduced the accumulation time distributions obtained from both stochastic simulations, even when the simulations violated some of the model’s underlying assumptions. We then demonstrated that the Moran process can be fitted directly to cross-sectional data, using summary statistics such as the fraction of cells exceeding different mutant thresholds. Applying global optimization and parametric bootstrapping, we showed that key parameters like mutation probability, selection advantage, and fraction of advantageous mutants, can be recovered with high accuracy and well-defined confidence intervals.

Importantly, we quantified how parameter uncertainty scales with sample size, demonstrating that confidence intervals derived from synthetic data provide a principled estimate of uncertainty when analysing real scRNAseq datasets. This is particularly relevant given that experimental studies often involve only a few thousand cells. The results further highlight that selection advantage is the most informative parameter for discriminating between competing hypotheses, as it directly reflects whether mutant accumulation is driven by drift or positive selection.

In summary, we have developed and validated a comprehensive framework for extracting quantitative information about mtDNA mutant dynamics from cross-sectional single-cell data. By integrating stochastic simulations, analysis of mutant diversity, Moran process modelling, and robust parameter inference, we provide a unified approach to study mtDNA deletion accumulation. This framework not only reconciles several previously conflicting observations but also offers concrete, testable predictions for experimental datasets. Application of this approach to large-scale single-cell transcriptomic resources is essential for testing these predictions in vivo. Accordingly, a companion manuscript applying the framework to experimental data from the Tabula Muris Senis project^34^ has been submitted separately (see also our preprint at^33^), providing a direct analysis of mtDNA mutant accumulation dynamics in ageing mouse tissues and extending the methodological results presented here.

## Methods

### Stochastic simulation of mitochondrial life-cycle

The stochastic simulations were implemented in Java and are based on an individual-based representation of mitochondrial DNA molecules within a single post-mitotic cell. Each simulation trajectory represents the lifetime dynamics of mtDNA populations under replication, degradation, and mutation, corresponding to the stochastic analogue of the deterministic ODE models described in the main text.

At any time point, the cellular mtDNA population is represented as a set of mtDNA “types”, where each type corresponds to a distinct mtDNA species (wild type or a specific deletion mutant) together with its current copy number. Each mutant type is assumed to be unique and does not further mutate. Each mtDNA type is associated with a replication rate, which for mutants may either equal the wild-type rate or take a higher value, reflecting a replication advantage.

Time is discretized in steps of one hour. At each step, two stochastic processes are applied, representing degradation and replication. Degradation is modelled as a binomial process, where for each mtDNA type the number of molecules lost in a time step is drawn from a binomial distribution with parameters given by the current copy number and a degradation probability derived from the specified half-life. Replication is likewise treated stochastically: for each mtDNA type, the expected number of replication events is determined based on the ODEs described in the main text. The actual number of replication events is then drawn from a binomial distribution. For wild-type mtDNA, each replication event can give rise to a deletion mutant with a specified mutation probability; newly arising mutants are assigned their replication rate at birth and are tracked as separate types thereafter. All degradation, replication, and mutation events are sampled using optimized binomial random number generators based on the Marsaglia–Tsang–Wang algorithm, as implemented in the Apache Commons RNG library.

Simulation output is recorded to a data file at fixed reporting intervals specified by the user (e.g. 30 days). At each reporting point, the current mtDNA composition is stored, including the copy numbers of all mtDNA types and their associated replication rates. A trajectory is considered “taken over” when the fraction of mutant mtDNA exceeds a predefined threshold.

### Using the Moran process to model accumulation times

To model the spread of mtDNA mutants, we employed the Moran process^28-30^, a classical stochastic model from population genetics that is well suited for describing how a new variant spreads in a finite population of constant size. In the Moran model, each step consists of two events, the random selection of one individual that will reproduce and another is randomly selected to die, so that the population size remains fixed. The probability that a particular type is chosen for reproduction depends on its relative fitness. As a result, a mutant with a selective advantage is slightly more likely to increase in number, whereas a neutral or disadvantaged mutant may still drift but is more prone to extinction. Over time, the system can only end in one of two absorbing states, the mutants either go extinct or they reach fixation, fully replacing the wild-type population.

We model a cell containing N (e.g. 1000) mtDNA molecules, starting with a single mutant and the remainder wild type. Types differ by a multiplicative replication advantage, with fitness values given by *f*_*α*_ = 1 + *s*_*α*_ (wild type s_α_ = 0). If the mutant carries a selection advantage, this bias alters the birth–death dynamics in its favour. The central quantity of interest is the distribution of times required for the mutant population to expand to dominance. Because the Moran process is a discrete-time Markov chain with well-defined transition probabilities between states {i = 0, 1, …, N}, it is possible to calculate this distribution exactly. We used a recursive first-step analysis:

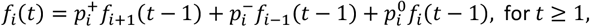

with boundary conditions:

- *f*_*N*_(0) = 1 (if we start fully mutant, fixation time is 0 with probability 1),
- *f*_*i*_(0) = 0 (cannot be fixed at time 0 if not already in state N).

The probability of fixation at time t from state *i* can be written in terms of the probabilities of moving to neighbouring states at the first step and then fixing in *t−1* steps from there. With suitable boundary conditions, the full distribution can be built iteratively. Alternative formulations, such as spectral decompositions of the transition matrix^35^, provide equivalent results and confirm that fixation time distributions are typically broad and right-skewed.

### Using the Moran process to calculate probability distributions

To obtain probability distributions rather than single stochastic trajectories, we write the forward master equation for the count state m = (m_1_,…,m_K_) of *K* mutant classes and wild-type 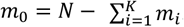. The only allowed transitions are m → m + e_γ_ − e_δ_, representing one birth of type γ and one death of type δ. The transition probability for a Moran step (from state **m**) involving one birth of type γ and one death of type δ is given by:

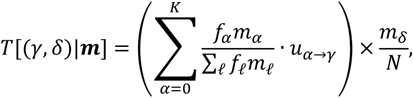

This yields a sparse, time-homogeneous Markov kernel on the feasible lattice {***m***: *m*_*i*_ ≥ 0, ∑_*i*_ *m*_*i*_ ≤ *N*}. Starting from a pure wild-type initial condition, we propagate the probability tensor P_t_(m) forward by one step with this kernel and sample it at the reporting times.

To map from real time to Moran steps, we use the organelle replication rate *r*_*b*_ = ln 2 /halfL, so that one ‘Moran step’ corresponds to one replication and degradation event in the cell. Thus over Δt days the expected number of steps is *Nr*_*b*_Δ*t*. Computationally, this enables efficient vectorized updates on the grid. This exact, mechanistic construction produces time-resolved probability distributions for multiple mutant classes with distinct selection advantages, parameterized in physical units.

For parameter estimation, we minimized the sum of squared differences between the Moran predictions and the empirical curves across all time points. For optimization we employed the Python library “fcmaes” (https://github.com/dietmarwo/fast-cma-es), specifically its Differential Evolution (DE) algorithm. DE is a population-based, gradient-free optimization method, well suited for nonlinear, multimodal optimization problems. To obtain parameter estimates together with statistically meaningful uncertainty measures, we employed parametric bootstrapping. Specifically, we generated 40 independent synthetic data sets using stochastic simulations of the Moran model with known parameters. Each synthetic data set was then analysed using the same parameter fitting pipeline. From the resulting empirical distributions of parameter estimates, 95 % confidence intervals (CI) were obtained as the 2.5th and 97.5th percentiles.

## Data and Code Availability

Data and computer code used for this work is available via our GitLab repository at https://gitlab.uni-rostock.de/IBIMA/MitoMutantAccTimes1.

## Acknowledgements

We would very much like to thank Dr. D. Wolz for his help with parameter fitting using fcmaes (https://github.com/dietmarwo/fast-cma-es) and for optimizing the speed of the calculation of mutant distributions based on the Moran process.

## Author Contributions

AK designed the overall concept of the manuscript and developed the mathematical simulations. AK and TK were involved in writing the text and all authors reviewed and approved the manuscript.

## Competing Interests

The authors declare that there are no conflicts of interest.

## Notes

### Competing Interest Statement

The authors have declared no competing interest.

### Summary of Updates

This revision contains an updated reference.

